# p53 modulates kinase inhibitor resistance and lineage plasticity in NF1-related MPNSTs

**DOI:** 10.1101/2023.01.18.523629

**Authors:** Jamie L. Grit, Lauren E. McGee, Elizabeth A. Tovar, Curt J. Essenburg, Emily Wolfrum, Ian Beddows, Kaitlin Williams, Rachael Sheridan, Josh Schipper, Marie Adams, Menusha Arumugam, Thomas Vander Woude, Sharavana Gurunathan, Jeffrey M. Field, Julia Wulfkuhle, Emanuel F. Petricoin, Carrie R. Graveel, Matthew R. Steensma

**Affiliations:** Department of Cell Biology, Van Andel Research Institute, Grand Rapids, MI 49503, USA; Bioinformatics & Biostatistics Core, Van Andel Research Institute, Grand Rapids, MI 49503, USA; Flow Cytometry Core, Van Andel Research Institute, Grand Rapids, MI 49503, USA; Genomics Core, Van Andel Research Institute, Grand Rapids, MI 49503, USA; Department of Pharmacology, University of Pennsylvania Perelman School of Medicine, Philadelphia, PA 19104, USA; Center for Applied Proteomics and Molecular Medicine, George Mason University, Manassas, VA 20110, USA; Helen DeVos Children’s Hospital, Spectrum Health System, Grand Rapids, MI 49503, USA; Michigan State University College of Human Medicine, Grand Rapids, MI 49503, USA

**Author notes:** Corresponding Authors: Carrie R. Graveel, PhD, Van Andel Research Institute, 333 Bostwick Ave. NE, Grand Rapids MI, 49503, Van Andel Research Institute, 333 Bostwick Ave. NE, Grand Rapids MI, 49503.

**Keywords:** NF1, MPNST, p53, resistance, plasticity

## Abstract

Malignant peripheral nerve sheath tumors (MPNSTs) are chemotherapy resistant sarcomas that are a leading cause of death in neurofibromatosis type 1 (NF1). Although NF1-related MPNSTs derive from neural crest cell origin, they also exhibit intratumoral heterogeneity. *TP53* mutations are associated with significantly decreased survival in MPNSTs, however the mechanisms underlying *TP53-*mediated therapy responses are unclear in the context of *NF1*-deficiency. We evaluated the role of two commonly altered genes, *MET* and *TP53*, in kinome reprograming and cellular differentiation in preclinical MPNST mouse models. We previously showed that *MET* amplification occurs early in human MPNST progression and that *Trp53* loss abrogated MET-addiction resulting in MET inhibitor resistance. Here we demonstrate a novel mechanism of therapy resistance whereby p53 alters MET stability, localization, and downstream signaling leading to kinome reprogramming and lineage plasticity. *Trp53* loss also resulted in a shift from RAS/ERK to AKT signaling and enhanced sensitivity to MEK and mTOR inhibition. In response to MET, MEK and mTOR inhibition, we observed broad and heterogeneous activation of key differentiation genes in *Trp53*-deficient lines suggesting *Trp53* loss also impacts lineage plasticity in MPNSTs. These results demonstrate the mechanisms by which p53 loss alters MET dependency and therapy resistance in MPNSTS through kinome reprogramming and phenotypic flexibility.

## Introduction

Malignant peripheral nerve sheath tumors (MPNSTs) are aggressive, chemoresistant sarcomas arising from Schwann cells that are the leading cause of death in patients with Neurofibromatosis Type 1 (NF1) (1). NF1 is an autosomal dominant tumor predisposition syndrome caused by inactivating mutations in the *NF1* gene (2–4). *NF1* is a tumor suppressor gene that encodes neurofibromin, a critical negative regulator of RAS (5). NF1-related MPNSTs exhibit deregulated RAS signaling caused by loss of heterozygosity of *NF1* along with additional tumor suppressor loss (*TP53, CDKN2A, SUZ12, PTEN*) and receptor tyrosine kinase (RTK) amplification (*MET, EGFR, PDGFR*) (6–12). As such, targeted therapies against RTKs and RAS effectors including MEK have been proposed as a treatment option for MPNSTs. Even with promising preclinical results, clinical trials featuring tyrosine kinase inhibitors have not been successful to date (13–16). Because of their aggressive clinical behavior, the 5 year survival rate remains only at 10-50% (17–20).

Although histologic and genomic MPNST subtypes have been described, these categories are not therapeutically relevant. The MPNST chemotherapy regimen has remained largely unchanged since the incorporation of doxorubicin in the 1980’s. The lack of actionable MPNST subtypes remains a major barrier to effective treatment, particularly given the vast differences in kinase signaling between histologically identical tumors (14,21–25). Additionally, MPNSTs are known to exhibit divergent states of differentiation leading to intratumoral heterogeneity. For example, MPNSTs can contain cellular regions comprised of malignant muscle, bone, fat, nerve and cartilage cells (26). It has long been suspected that the differentiation states of various MPNST histologic subtypes contribute to therapy resistance, but confirmatory data is lacking. Thus the identification of predictive biomarkers for MPNSTs continues to be an area of intense study (27). Activation of the PI3K/AKT/mTOR pathway is a common event, occurring in approximately 50% of MPNSTs, and is associated with poor prognosis (28). Additionally, 25-60% of MPNSTs are p53-deficient (9,10,29), which is associated with significantly diminished survival (29–31) and poor response to neo-adjuvant chemotherapy (32). Defining the role of these pathways in therapeutic response is critical to predicting effective targeted therapies and predictive biomarkers for future MPNST trials.

Consistent with these clinical observations, we previously found that a *Trp53*-deficient mouse model of NF1-related MPNST exhibited faster tumor growth and was more resistant to chemotherapy compared to *Trp53*-intact models. Interestingly, the *Trp53*-deficient model was also less sensitive to single agent and combination MEK and MET inhibition despite sustained repression of ERK phosphorylation. *Trp53*-deficient tumors also exhibited unusually high AKT activation both at baseline and in response to targeted therapy (25,33). In this study, we aimed to define the role of p53 in regulating kinase signaling, targeted therapy response, and cellular differentiation in NF1-related MPNSTs. Transcriptomic and phospho-proteomic analysis revealed multiple mechanisms of resistance, including deregulation of MET stability and localization, deregulated PI3K/AKT/mTOR signaling, and altered lineage plasticity. In contrast to these results, we found that p53 loss actually increased sensitivity to mTOR inhibition, which was associated with broad and persistent kinome activation. Excitingly, combined mTOR and MEK inhibition reversed clonal selection for p53-deficiency and was the most effective drug combination in all models, regardless of p53 status. Lineage plasticity, as defined by transcriptional profiling, was closely linked to tyrosine kinase inhibitor resistance. Collectively, these data indicate that p53 acts as a master regulator of tyrosine kinase signaling and mediates oncogene-addiction and cell fate in MPNSTs.

## Methods

### Cell Culture and Drugs

Mouse-derived MPNST cell lines described previously (33) were cultured at 37 °C in a humidified atmosphere in 5% CO_2_ in low pH DMEM (Thermo Fisher Scientific) supplemented with 10% FBS (Corning, lot #35070165) and 1% Penicillin Streptomycin (Thermo Fisher Scientific), unless otherwise indicated. Cell lines were verified to be free of *Mycoplasma* contamination every 6 months by PCR (ATCC). Human-derived MPNST cell lines were cultured as described previously (34). Capmatinib (Novartis), trametinib (Novartis), everolimus (Selleckchem), and afuresertib (Selleckchem) solutions were prepared in DMSO. For human cell line IC_50_ experiments, drugs were purchased as 10 mM stock solutions (Selleckchem) and handled as previously described (34).

### Viability and Dose Response

For cell viability experiments, cells were allowed to adhere and then treated with DMSO or the indicated drug dose. After 72 hours, cells were trypsinized (Thermo Fisher Scientific) and trypan blue negative cells were counted using a TC20 Automated Cell Counter (Bio-Rad) to determine percent viability relative to vehicle treatment. Pairwise comparisons of beta regressions were done using the betareg (v 3.1-3) and emmeans (v 1.4.5) packages and plotted using the ggplot2 (v 3.3.0) package in R (v 3.6.3). Emmeans and ggplot2 were also used for comparison of doubling times. For human MPNST cell lines, the IC_50_ was determined as previously described (34). Briefly, cells were allowed to adhere and then treated with DMSO or the indicated serially diluted drug (4.6 nM – 10 μM) for 72 hours, cell viability was measured by ATPlite Luminescence Assay (PerkinElmer), and IC_50_ was calculated using GraphPad Prism 7. The rcorr function in the Hmisc (v4.4-1) package was used to generate spearman correlations of IC_50_. The ggplot2 (v 3.3.0) and ggpubr (v 0.2.5) packages were used to plot an IC_50_ matrix and correlogram, respectively, using R (v 3.6.3). For dose combination matrices, 2500 cells/well were plated in a 96 well plate, allowed to adhere, and then treated with the indicated serially diluted drug or vehicle for 72 hours. Cell viability was measured using CellTiter 96 Aqueous One Solution Cell Proliferation Assay (MTS) (Promega) following the manufacture’s protocol. Absorbance was read using Synergy Neo microplate reader (BioTek) and normalized to cell number using a standard curve generated by GraphPad Prism 7. Mean percent inhibitions relative to vehicle controls were plotted in dose response matrices using the synergyfinder package (v 2.0.12) in R (v 3.6.3).

### Reverse Phase Protein Array

For cell culture experiments, cells were seeded in a 6-well plate, allowed to adhere overnight, and then treated in replicates of 6 with DMSO or the indicated drug or ligand dose for 2 or 48 hours. To harvest samples, the cells were washed thoroughly 3 times with ice cold PBS, and plates were immediately snap frozen on dry ice to preserve the integrity of the phosphoproteome. For mouse tumorgrafts, we analyzed data, where some of this data was used in a previous study (25). Briefly, immediately following the euthanasia of tumor-bearing mice, 15–25 mg portions of each tumor were transplanted into the flank of NSG-SCID mice using a 10-gauge trochar. When the tumor volume reached approximately 150 mm^3^, mice were euthanized, and tumors were immediately harvested and snap-frozen in liquid nitrogen within 20 min upon surgical resection to preserve the integrity of the phosphoproteome. Six tumors were assessed for each genotype. All animal experimentation in this study was approved by the Van Andel Institute’s Internal Animal Care and Use Committee (XPA-19-04-001). Specimens were then embedded in an optimal cutting temperature compound (Sakura Finetek, Torrance, CA, USA), cut into 8 μm cryo-sections, mounted on uncharged glass slides, and stored at − 80 °C until use. Each slide was fixed in 70% ethanol (Sigma Aldrich, Darmstadt, Germany), washed in deionized water, stained with hematoxylin (Sigma Aldrich, Darmstadt, Germany) and blued in Scott’s Tap Water substitute (Electron Microscopy Sciences), and dehydrated through an ethanol gradient (70%, 95%, and 100%) and xylene (Sigma Aldrich, Darmstadt, Germany). In order to prevent protein degradation, complete protease inhibitor cocktail tablets (Roche Applied Science, Basel, Switzerland) were added to the ethanol, water, hematoxylin, and Scott’s Tap Water substitute (35). Cells were lysed in a 1:1 solution of 2× Tris-Glycine SDS Sample buffer (Invitrogen Life Technologies, Carlsbad, CA, USA) and Tissue Protein Extraction Reagent (Pierce, Waltham, MA, USA) supplemented with 2.5% of 2-mercaptoethanol (Sigma Aldrich, Darmstadt, Germany). Cell lysates were boiled for 8 min and stored at − 80°C. Reverse phase protein microarray construction and immunostaining was performed as previously described (25) A total of 98 protein sites passed quality control metrics and were used for analysis. Differential expression was performed using R package “limma” (36) and R (v 3.6.0. For differential activation analysis, P-values for delta-delta differences in limma fold changes were generated in R (v 3.6.3) using Wald tests and adjusted using the BH method. Adjusted P-values of less than 0.05 were considered significant. For tumors, the pairwise difference between genotypes in mean centered phospho-site expression (Z score) was assessed using a linear mixed-effects model with a random intercept for each phospho-site using the lme4 (v 1.1-23) and emmeans (v 1.4.5) packages in R with the selected phospho-sites. Pairwise differences between genotypes in log transformed expression were also assessed for each individual phospho-site using emmeans. A comprehensive analysis including all 98 tumorgraft phospho-sites was previously published (25) Balloon plots were created from the limma fold changes (cell lines) or z-scores (tumors) in R (v 3.6.3) using the ggballoonplot function in the ggpubr package (v 0.2.5). Labeled fold change, differential activation, waterfall, and site-specific expression plots were created using ggplot2 (v 3.3.0) in R (v 3.6.3).

### Lentiviral transduction

To generate the NF1-MET;sgP53 cell line, the pLentiCRISPRv2 vector (Addgene #52961) (37) was digested with BsmBI and dephosphorylated with CIP (New England BioLabs) following the manufactures instructions and then purified with using Qiaquick Gel Extraction kit (Qiagen). The oligonucleotides 5’-cac cgA GCC AAG TCT GTT ATG TGC A-3’ and 5’-aaa cTG CAC ATA ACA GAC TTG GCT c-3’ were annealed and then ligated into the vector using T4 PNK and T4 ligase (Thermo Fisher Scientific). One Shot competent cells (Thermo Fisher Scientific) were transformed and candidate colonies were verified by diagnostic digestion. Lentivirus was produced by lipofectamine transfection (Thermo Fisher Scientific) of HLA 293 cells kindly provided by Dr. Bart Williams with either the pLentiCRISPRv2;sgP53 plasmid or the pLentiCRISPRv2 plasmid (as an empty vector control), along with the envelop and packaging plasmids pCMV-VSV-G (Addgene #8454) (38) and psPAX2 (Addgene #12260). NF1-MET cells were infected with 500 μL of 0.45 μm filtered virus with polybrene (EMD Millipore), and 48 hours post infection transduced cells were selected for by 2 μg/mL puromycin (Thermo Fisher Scientific) treatment. To generate H2B GFP and RFP labeled cells, lentivirus was produced by lipofectamine transfection of Phoenix 293 cells kindly provided by Dr. Bart Williams with either LV-GFP (Addgene #25999) (39) or pHIV-H2BmRFP (Addgene #18982) (40) plasmids, along with pCMV-VSV-G and psPAX2. Filtered lentivirus was concentrated using Lenti-X Concentrator (Takara). NF1-MET;sgEmptyVector and NF1-MET;sgP53 cells were infected with GFP or RFP lentivirus to create NF1-MET-GFP and NF1-MET;sgP53-RFP cell lines, respectively.

### Competition Assay

For clonal competition assays, single cell suspensions of NF1-MET-GFP and NF1-MET;sgP53-RFP cells were prepared in sorting buffer (HBSS without Ca^2+^, Mg^2+^, and Phenol Red (Thermo Fisher Scientific) with 25 mM HEPES, 2 mM EDTA, 2% FBS, and 10 mg/mL DAPI added). 7,500 GFP+ and 7,500 RFP+ single live cells were sorted into each well of a 24 well culture plate containing low pH DMEM with 10% FBS using a MoFlo Astrios cell sorter with Summit v6.3 software (Beckman Coulter). Sorting was performed at 25psi using a 100um nozzle, with Purify for the abort mode and a drop envelope set to 1-2 drops. Cells were selected using SSC vs FSC, and single cells using both SSC area vs height and area vs width. Live cells (DAPI negative) were identified using the 355-448/59 channel, and GFP and RFP signals were detected using the 488-510/20 and 561-614/20 channels, respectively.

After cells adhered, they were treated with DMSO or the indicated drug dose and cocultured for the indicated timepoints. The cocultured cells were analyzed to determine the percentage of single, live cells that were either GFP+ or RFP+ using a CytoFLEX S flow cytometer with CytExpert v2.4 software (Beckman Coulter). Analysis was performed using FlowJo v10.7 software (BD Life Sciences). Single cells were identified using SSC vs FSC, followed by SSC area vs height. Live cells (DAPI negative) were identified using the 405-450/45 channel, and GFP and RFP signals were detected using the 488-525/40 and 561-610/20 channels, respectively. Pairwise comparisons of beta regressions were done using the betareg (v 3.1-3) and emmeans (v 1.4.5) packages and plotted using the ggplot2 (v 3.3.0) package in R (v 3.6.3).

### Western Blotting

Cells were grown overnight followed by serum starving overnight for a final confluency of 90%, and then treated with drug and/or stimulated with 10% serum or 100 ng/mL HGF as indicated. Cell lysate collection and immunoblotting were done as previously described (33). Primary antibodies were purchased from Cell Signaling: p53 (#2524), phospho-MET Y1234/1235 (#3077), MET (#3127), phospho-AKT S473 (#9271), phospho-ERK T202/Y204 (#9101), phospho-S6 S235/236 (#4858), β-Actin (#3700).

### RT-qPCR

RNA was isolated using RNeasy Mini Kit and RNase-Free DNase set (Qiagen) following the manufactures protocol. cDNA was synthesized using SuperScript II Reverse Transcriptase with Oligo (dT) 12-18mer Primers (Thermo Fisher Scientific) using Tetrad 2 (BioRad) following the manufactures protocols. qPCR was done on Step One Plus (Applied Biosystems) using Fast Start Universal SYBR Green Master Mix (Roche) with the following primers: Cdkn1a: Forward 5’-CTT GCA CTC TGG TGT CTG-3’ Reverse 5’-CTT GGA GTG ATA GAA ATC TGT CA-3’; Tubb5: Forward 5’-ATG CCA TGT TCA TCG CTT AT-3’ Reverse 5’-TTG TTC GGT ACC TAC ATT GG-3’. Relative expression was calculated using the ddCt method and plotted using GraphPad Prism 7.

### Mouse Xenograft Models

Six- to eight-week-old female NSG-SCID mice were injected subcutaneously with one million cells of either NF1-MET or NF1-MET;sgP53 (10 mice/cell line). Tumors were measured twice weekly with calipers. Once tumors reached approximately 150 mm^3^, 5 mice from each cell line group were treated with either 30 mg/kg capmatinib or vehicle (0.5% methycellulose) twice daily for 15 days or until tumors reach 2500 mm^3^. A second experiment was performed exactly the same way, for a final total of 10 mice in each cell line and treatment group.

A linear mixed-effects model was used to determine if there were significant differences between the results of experiment one and experiment two. The models were stratified by treatment group. Data was square root transformed before it was entered into the model. The model used a 3-way interaction for timepoint, cell-line, and experiment and also included a random intercept for tumor ID. A linear mixed-effects model was used to run the analysis for the combination analysis. Data was square root transformed before it was entered into the model. This was a linear mixed effects model with a 4-way interaction between timepoint, experiment, treatment group, and cell-line. A random intercept for tumor-id was also included. All statistical analyses were run using R (v. 3.6.0) and all linear regression models were run using the lme4 package. Emmeans was used for the contrasts/comparisons that were run on the regression output. All p-values that were obtained through the emmeans output were automatically multiple-testing corrected via the Tukey method.

### Immunofluorescence

Cells were plated at 50,000 cells per well on glass coverslips (Fisherbrand) in a 24 well plate and allowed to attach overnight. The next day cells were serum-starved for 16 hours. The third day cells were treated with 100 ng/mL HGF for 5 minutes at 37°C, washed three times with phosphate buffered saline + 0.5% tween 20 (PBST), and fixed with 4% formaldehyde in PBS for 15 minutes. Cells were permeabilized with ice-cold methanol for 3 minutes at -20°C. Samples were blocked (5% normal goat serum, 1% BSA, 0.3% Triton X-100 in PBS) for one hour at room temperature in a humidity chamber. Antibodies against phospho-MET (Y1230/1234/1235; Abcam #5662) were diluted 1:100 in antibody dilution buffer (1% BSA and 0.3% triton-x 100 in PBS) and incubated with the cells overnight at 4°C in a humidity chamber. Primary antibodies were detected using Alexa Flour secondary antibody goat-against-rabbit 594 (Life Technologies #8889) at 1:500 and incubated for 40 minutes at room temperature. Cell nuclei were stained with DAPI and samples were mounted with ProLong Gold antifade reagent (Life Technologies). For imaging, at three regions of interest were imaged per sample using a 60x Plan Apo VC oil immersion objective with 1.4 NA on a Nikon A1 plus-RSi laser scanning confocal microscope (Nikon Elements software). Image resolution was 1024×1024 of z-slices covering the entirety of cell thickness. PMT levels were set using controls with 403 and 561 solid-state lasers. FIJI was used to generate maximum intensity projection TIFF images.

### RNA Sequencing

Cells were plated in a 6-well plate, grown in 10% FBS overnight, and then treated in duplicate with 100 nM capmatinib or DMSO. RNA was isolated using RNeasy Mini Kit and RNase-Free DNase set (Qiagen) following the manufactures protocol. Libraries were prepared by the Van Andel Genomics Core from 500 ng of total RNA using the KAPA mRNA Hyperprep kit (v4.17) (Kapa Biosystems, Wilmington, MA USA). RNA was sheared to 300-400 bp. Prior to PCR amplification, cDNA fragments were ligated to IDT for Illumina TruSeq UD Indexed adapters (Illumina Inc, San Diego CA, USA). Quality and quantity of the finished libraries were assessed using a combination of Agilent DNA High Sensitivity chip (Agilent Technologies, Inc.), QuantiFluor® dsDNA System (Promega Corp., Madison, WI, USA), and Kapa Illumina Library Quantification qPCR assays (Kapa Biosystems). Individually indexed libraries were pooled and 50 bp, paired end sequencing was performed on an Illumina NovaSeq6000 sequencer using an S2, 100 bp sequencing kit (Illumina Inc., San Diego, CA, USA) to a minimum raw depth of 41.4 M reads with an average of 47.2 M reads per sample. Base calling was done by Illumina RTA3 and output of NCS was demultiplexed and converted to FastQ format with Illumina Bcl2fastq v1.9.0. Sequencing adapters were trimmed using Trimgalore v0.4.2 (http://www.bioinformatics.babraham.ac.uk/projects/trim_galore/). Bases with a quality score less than 20 were also removed from the ends of reads. Trimmed data were quality controlled with FastQC v0.11.7 (41) and then mapped with STAR v2.5.2b to the mm10 genome using the default settings (42). Raw gene counts (minimum of 30M counts per sample and a mean of 34.92) generated by STAR were imported into R v3.6.0. Genes with less than 10 counts in a minimum of two samples were immediately removed from all further analysis; this minimizes multiple testing adjustments and removes low-expressed genes unlikely to be biologically meaningful. For all differential expression contrasts using a subset of the data, the count data were further filtered so that, for the samples being used in the contrast, a minimum of two samples have >0 counts. A quasi-likelihood negative binomial generalized log-linear model was then fit to the filtered count data using the weighted trimmed mean of M-values to normalize for library size and composition biases (43). P-values were generated using empirical Bayes quasi-likelihood F-tests, and then adjusted using the BH method; adjusted P-values less than 0.05 were considered significant. Gene ontology enrichment analyses were done using the clusterProfiler R package v3.14.3 using the function ‘goseq’ with KEGG ontologies. Heatmaps were generated from library-size normalized counts centered across genes (z-scores) using the pheatmap package v1.0.12. Intersects were determined and plotted using the R package UpSetR v1.4.0 using the ‘upset’ function (44).

## Results

### p53-deficiency is associated with MET inhibitor resistance in MPNSTs

Previously, we compared response to MET and MEK inhibition in tumor xenografts derived from MPNSTs of genetically engineered mouse models of NF1, including a *Met*-amplified, *Trp53*-wildtype model (NF1-MET) and a *Trp53* deficient model (NF1-P53). NF1-P53 MPNSTs were less sensitive to both MET and MEK inhibition *in vivo* (33). To confirm the MPNST cell lines maintained their drug sensitivity phenotypes *in vitro*, we generated tumor cell line isolates and performed cell viability analysis after 72 hours of drug treatment. (Fig. 1A). These results validated that the NF1-P53 cell line was resistant to not only single agent MET (capmatinib) and MEK (trametinib) inhibition, but also was resistant to combined MET and MEK inhibition (Fig. 1A). A targeted analysis of the RAS/ERK and PI3K/AKT pathways with reverse phase protein arrays (RPPA) was used to identify both immediate (2 hours) and adaptive (48 hours) kinome responses to MET and MEK inhibition. After controlling for drug treatment and exposure time, several PI3K effectors (including AKT and S6) were significantly activated in the NF1-P53 cells, suggesting PI3K/AKT pathway activation may promote resistance to MET and MEK inhibition (Fig. 1B, Supplementary Fig. 1).

**Figure 1.**
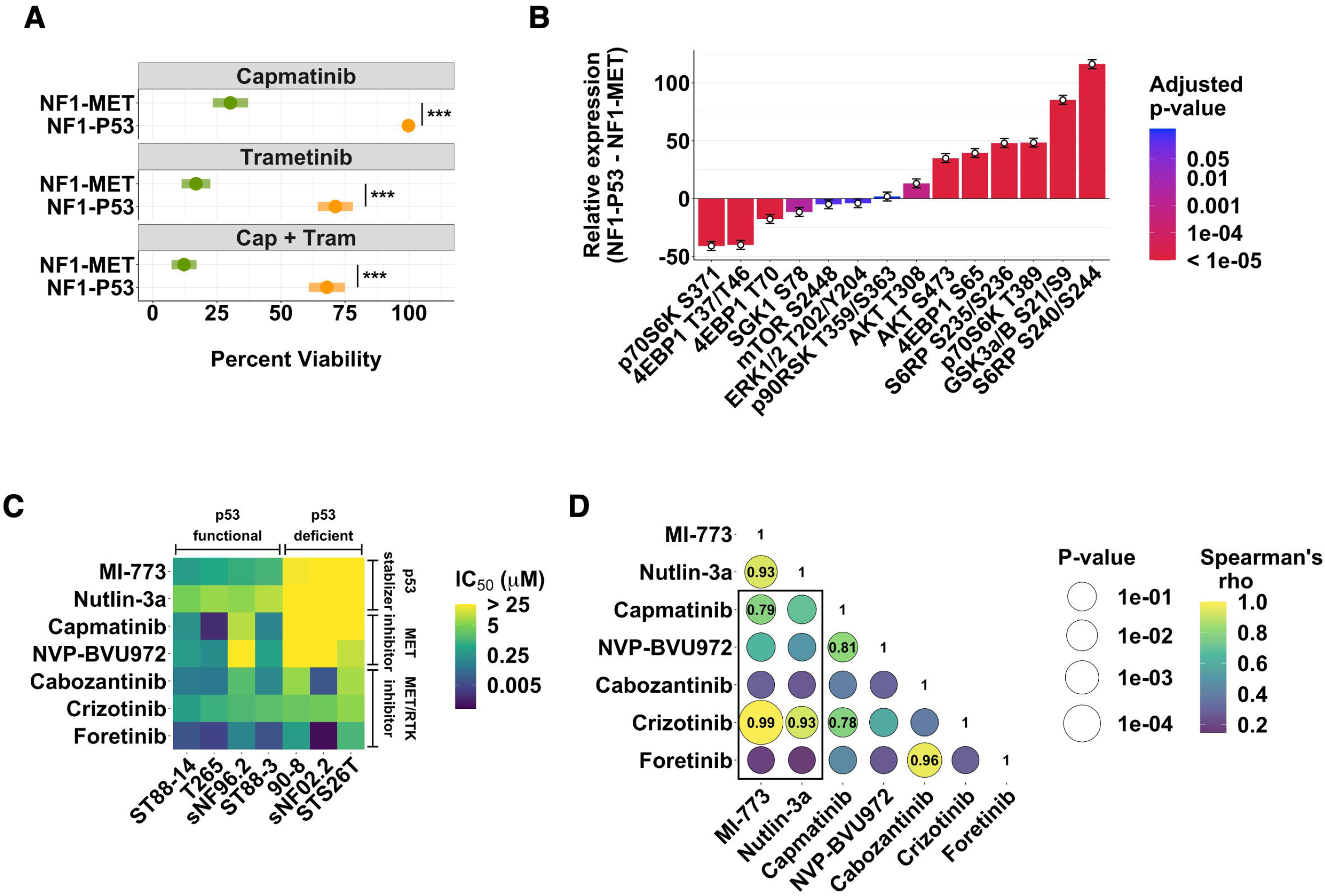
p53 deficiency is associated with MET inhibitor resistance in MPNSTs. (A) Percent viability of NF1-MET and NF1-P53 cells after 72 hours of capmatinib (100 nM), trametinib (40 nM) or combination (capmatinib 100 nM, trametinib 40 nM) treatment. (B) Change in phospho-site activation of MET effectors in NF1-P53 cells relative to NF1-MET cells upon capmatinib (100 nM), trametinib (100 nM), or combination (capmatinib 100 nM, trametinib 100 nM) treatment for 2 and 48 hours. See also Supplemental Figure 1. (C) IC_50_ of p53 stabilizing drugs and MET inhibitors against a panel of human MPNST cell lines. (D) Spearman’s correlations (color) and significance (size) between the IC_50_ of the drugs in panel C. The rho value of correlations with a p-value < 0.5 are indicated in their respective bubble. The black box indicates the correlations between p53 stabilizing drugs and MET inhibitors. *** p < 0.001

To determine if p53 also regulates sensitivity to MET inhibition in human MPNSTs, we screened a panel of 6 NF1-related and 1 sporadic (STS26T) MPNST cell lines for functional p53 based on sensitivity to the p53 stabilizing drugs, MI-773 and nutlin-3a. These p53 stabilizers inhibit MDM2-mediated p53 degradation and are selectively active in p53 wild-type cells (45). Based on the IC_50_ of these drugs, we classified four cell lines as being “p53-functional” and three cell lines as being “p53-deficient” (Fig. 1C). In addition, we screened the cell lines for sensitivity to of a panel of MET inhibitors. All but one of the p53-intact cell lines were sensitive to MET inhibition while, the p53-deficient cells were profoundly resistant to MET inhibition (Fig. 1C). The IC_50_ between several of the p53 stabilizers and MET inhibitors significantly correlated, similar to drugs within the same class (Fig. 1D), indicating a critical role for p53 in regulating MET-dependency.

### p53 loss drives MET inhibitor resistance in MPNSTs

To evaluate the impact of p53 loss of function in the context of MET-addiction, we used CRISPR-Cas9-mediated knockout of p53 in NF1-MET cells to create the NF1-MET;sgP53 line. p53 protein levels as well as the p53 target gene, p21, were diminished in NF1-MET;sgP53 cells (Supplemental Fig. 2A-B). Cell viability analysis demonstrated that NF1-MET;sgP53 cells were significantly less sensitive to MET inhibition than NF1-MET cells; however, combined MET-MEK inhibition restored drug sensitivity (Fig. 2A). We evaluated the impact of p53 on HGF-induced MET signaling and observed augmented MET signaling in NF1-MET;sgP53 cells, yet capmatinib inhibited HGF-dependent ERK and AKT activation regardless of p53 status (Fig. 2B). In contrast, in FBS-stimulated cells pERK and pAKT were minimally reduced with capmatinib treatment in NF1-MET;sgP53 cells suggesting capmatinib resistance may be partially mediated by parallel pathway activation that converges on ERK and AKT (Fig. 2C).

**Figure 2.**
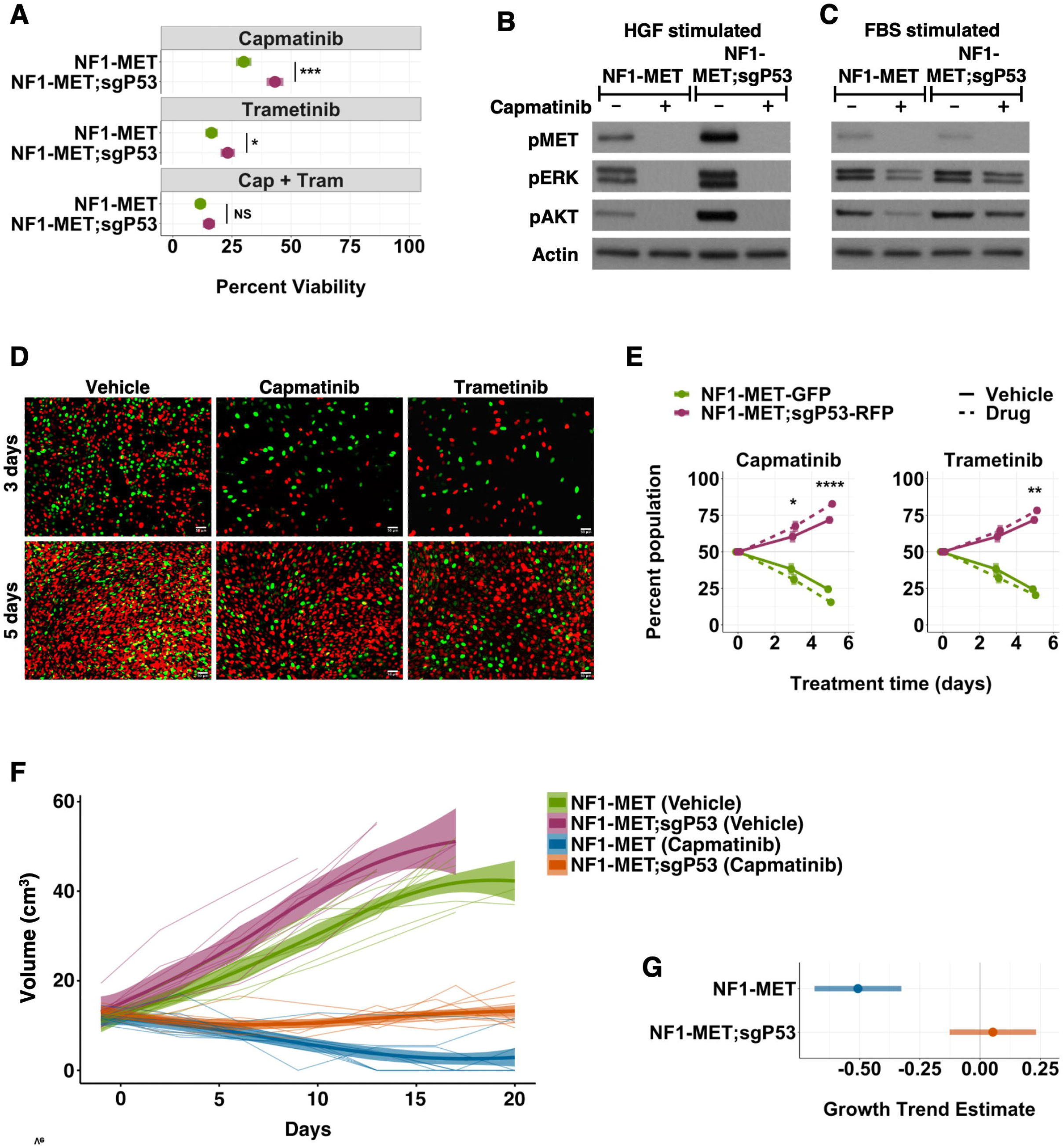
p53 loss drives MET inhibitor resistance in MPNSTs. (A) Percent viability of NF1-MET and NF1-MET;sgP53 cells after 72 hours of capmatinib (100 nM), trametinib (40 nM) or combination (capmatinib 100 nM, trametinib 40 nM) treatment. (B-C) Western blot of NF1-MET and NF1-MET;sgP53 cells treated with capmatinib (100 nM) for 2 hours and stimulated with HGF (B) or 10% FBS (C) for 15 minutes. Images (D) and flow cytometry analysis (E) of GFP labeled NF1-MET and RFP labeled NF1-MET;sgP53 cells after 3 and 5 days of treatment with vehicle (DMSO), capmatinib (100 nM), or trametinib (40 nM). (F) Individual tumor growth curves, LOESS curves, and 95% ribbons for vehicle or capmatinib (30mg/kg BID) treated NF1-MET and NF1-MET;sgP53 xenografts. (G) Pairwise comparison of growth trend estimates in capmatinib treated NF1-MET (p-value = 0.0083) and NF1-MET;sgP53 (p-value = 0.089) tumors to O (grey line). * p < 0.05, ** p < 0.01, *** p < 0.001, **** p < 0.0001

Resistant clonal populations within heterogeneous MPNSTs may explain the clinical failures of targeted kinase inhibitors in MPNSTs (46). To determine if p53 is a key driver of clonal selection and drug resistance in MPNSTs, we performed a clonal competition assay using labeled isogenic NF1-MET-GFP and NF1-MET;sgP53-RFP cell lines. Notably, when cultured separately, NF1-MET;sgP53 cells do not have a significant proliferative advantage compared to NF1-MET cells (Supplementary Fig. 2C). In contrast, when cocultured NF1-MET;sgP53-RFP cells had a strong growth advantage, which was significantly enhanced by both MET or MEK inhibition (Fig. 2D-E). After 5 days of capmatinib treatment, NF1-MET;sgP53-RFP cells comprised 83% of the culture (Fig. 2D-E), indicating that p53 loss drives strong clonal selection with MET inhibition.

To evaluate p53 loss and capmatinib sensitivity *in vivo*, we treated orthotopic MPNST xenografts with capmatinib. Tumor growth rate was significantly increased in the NF1-MET;sgP53 tumors compared to the parental tumors (p-value = 4.7e-4) (Fig. 2F; Supplementary Fig. 2D-E). Capmatinib strongly inhibited tumor growth in both the NF1-MET and NF1-MET;sgP53 models (Fig. 2F; Supplementary Fig. 2D), however only the NF1-MET tumors regressed on treatment (p-value = 0.0083), while the NF-MET;sgP53 tumors remained stable throughout treatment, with a subset of tumors actually showing increased growth (Fig. 2F-G; Supplementary Fig. 2D). These data confirm our *in vitro* findings and further demonstrate that p53 loss promotes MET inhibitor resistance.

### Met inhibition induces p53-dependent lineage plasticity

To understand how loss of p53 drives resistance we used RNA-seq to examine capmatinib-induced transcriptional changes in NF1-MET and NF1-MET;sgP53 cells. Unsupervised hierarchical clustering identified strong clustering by treatment followed by genotype, with p53 status modifying expression of some gene subsets (Fig. 3A). GO term enrichment analysis of genes upregulated by MET inhibition identified biological processes related to positive regulation of actin organization, cell adhesion, collagen deposition/ossification, and muscle differentiation (Fig. 3B, Supplementary Fig. 3A). We next examined genes that were downregulated in capmatinib treated NF1-MET;sgP53 cells and identified biological processes related to bone and kidney development (Fig. 3C, Supplementary Fig. 3B). Together, these data suggest that response to MET inhibition may be partially mediated through the induction of linage plasticity pathways in MPNSTs and that p53 loss disrupts this process to promote drug resistance. As differentiation and cell cycle arrest are coupled, we next examined whether p53 loss altered expression of known p53 target genes that promote cell cycle arrest. Indeed, expression of the cell cycle regulators *Cdkn1a* and *Zmat3* were lost in the NF1-MET;sgP53 cells, while expression of genes involved in senescence or apoptosis were unchanged (Supplementary Fig. 3C).

**Figure 3.**
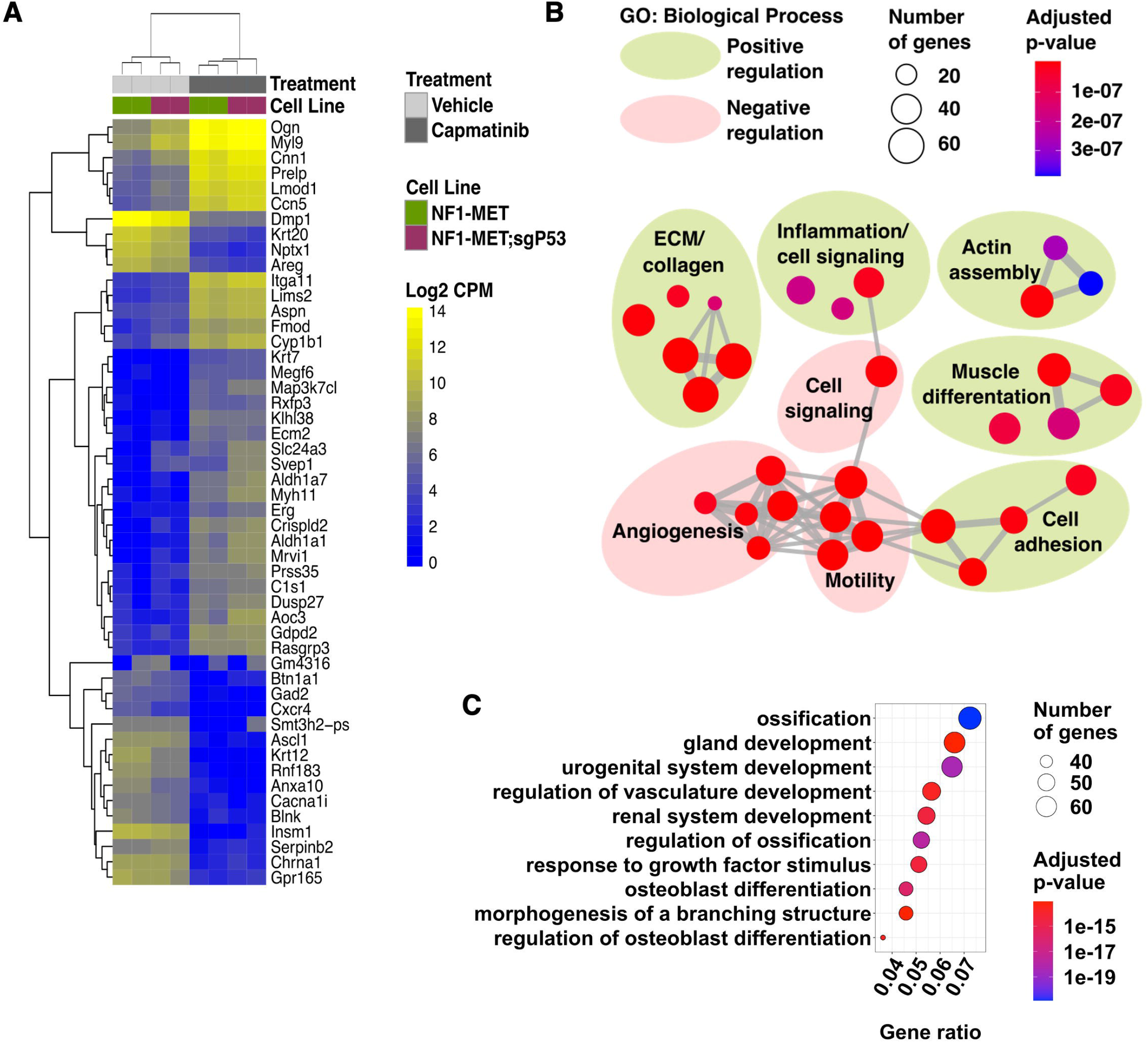
Met inhibition induces p53-dependent lineage plasticity. (A) Unsupervised hierarchical clustering of top 50 differentially expressed genes in NF1-MET and NF1-MET;sgP53 cells treated with capmatinib (100 nM) for 24 hours). (B) GO term enrichment analysis of capmatinib-induced genes. The top 30 most significantly enriched biological process terms (by adjusted p-value) are shown. Connecting grey lines represent relatedness of the pathways, while dot size indicates the number of genes differentially expressed in the pathway. See Supplemental Figure 3A for individual terms. (C) Top 10 most significantly enriched biological process GO terms derived from genes that were decreased in capmatinib treated NF1-MET;sgP53 cells. See Supplemental Figure 3B for expanded top 30 terms.

### p53 regulates MET stability and localization

RTK expression, activation, and recycling are tightly regulated processes that ensure RTK regulation in normal physiological conditions. Deregulation of this cycle of RTK activation and recycling is often observed in cancers, such as the MET exon 14 deletions observed in lung cancers (47). To determine whether RTK spaciotemporal regulation promoted the enhanced MET signaling observed in p53-deficient MPNST cells (Figs. 1B & 2B-C), we measured the kinetics of MET activation and turnover to increase effector activation. In NF1-MET cells an immediate, and expected, increase in MET activation was observed within 5 minutes of HGF treatment that quickly diminished along with total MET levels over time (Fig. 4A). In contrast, HGF-treatment induced a drastic pMET increase that remained elevated for 60 minutes in NF1-MET;sgP53 cells. Both phosphorylated and total MET were persistently elevated in the NF1-MET;sgP53 cells, which corresponded to increased and prolonged activation of both ERK and AKT (Fig. 4A). Activation of the mTOR effector S6 was similar between the two cell lines.

**Figure 4.**
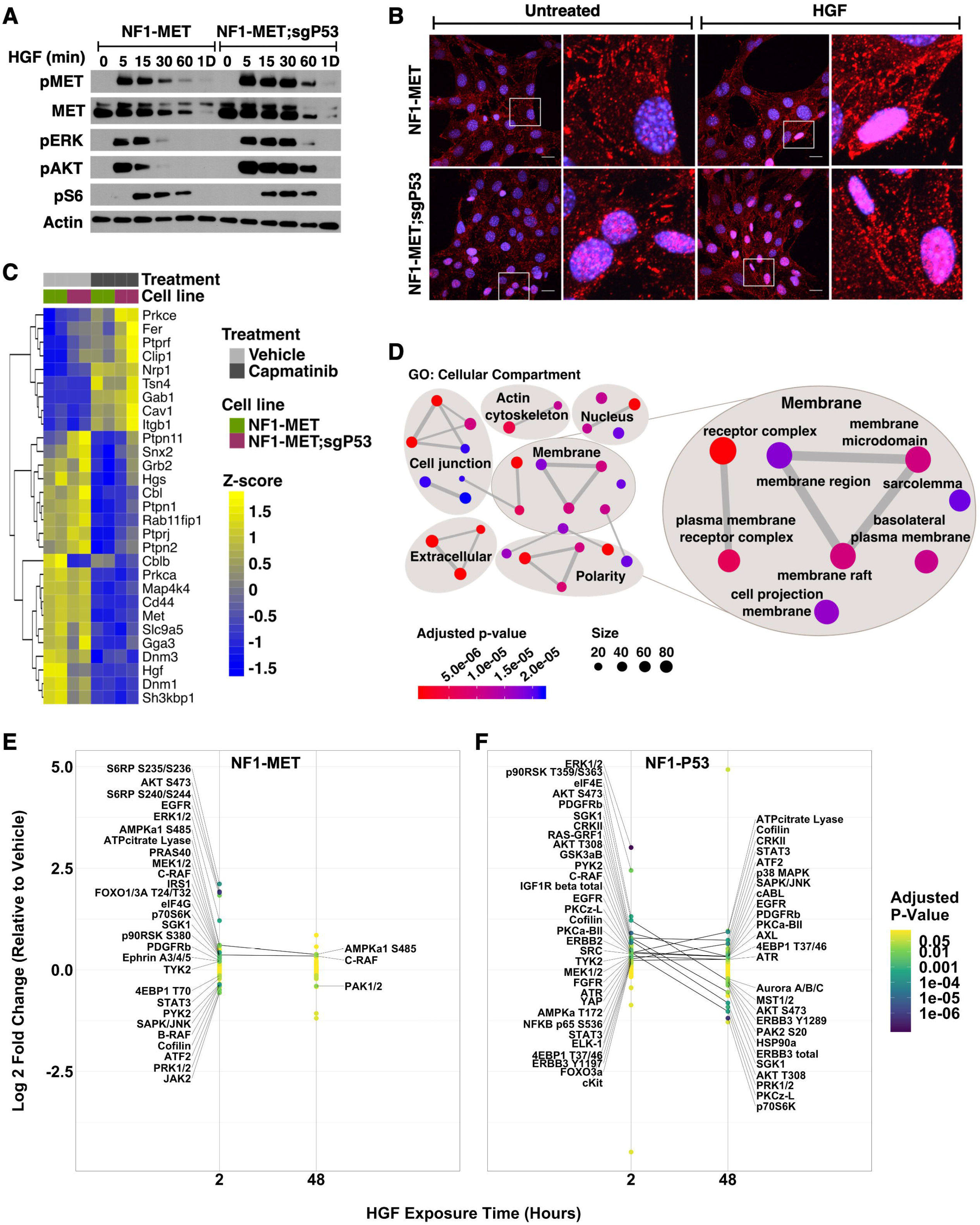
p53 regulates MET stability, localization, and effector signaling. (A) Time course western blot of NF1-MET and NF1-MET;sgP53 cells stimulated with HGF for 5 minutes to 1 day. (B) Representative images of phospho-MET localization (red) after 5 minutes of HGF stimulation. (C) Expression of genes associated with MET stability and localization after 24 hours of vehicle (DMSO) or capmatinib (100 nM) treatment. (D) GO analysis of Cellular Compartment terms downregulated in the NF1-MET;sgP53 cell line compared to the parental NF1-MET line. See Supplemental Figure 3D for individual terms. Change in phospho-protein expression in NF1-MET (E) and NF1-P53 (F) cells after HGF treatment. Significantly increased or decreased proteins at each time point are labeled and proteins that are significant at both the 2 and 48 hour time points are connected. See Table S1 for phosphosites. Color indicates P-value.

To examine whether p53 loss also alters MET subcellular localization, we performed immunostaining of MET after HGF-treatment. In normal conditions, HGF treatment results in MET localization and activation at the plasma membrane; however MET can also be internalized to the nucleus, where it’s function is incompletely understood (48). At baseline, pMET was localized to the cytoplasm in the NF1-MET cells, whereas we observed both nuclear and cytoplasmic staining in NF1-MET;sgP53 cells. Interestingly, HGF dramatically increased nuclear MET localization in the p53-deficient NF1-MET;sgP53 cells, while treatment induced nuclear MET localization only in a small percentage of NF1-MET cells (Fig. 4B). In several cancers, nuclear MET is associated with drug resistance and poor prognosis (49–53)). These results indicate that p53 loss induces nuclear MET localization, promoting tumor aggressive phenotypes in MPNST cells.

To understand how loss of p53 promotes increased stability and nuclear localization of MET, we used RNA-seq to examine the expression of genes involved in MET activation and turnover. Capmatinib treatment induced sweeping compensatory expression changes in both cell lines (Fig. 4C). In NF1-MET;sgP53 cells, expression of genes critical for MET degradation, *Sh3kpb1* and *Cblb*, (54–56) were significantly downregulated compared to the parental cell line. Conversely, *Prkce* expression, which is required for nuclear MET translocation (57), was significantly upregulated in the NF1-MET;sgP53 cells in the presence of capmatinib. GO cellular compartment enrichment analysis revealed that p53 loss promoted significant downregulation in plasma membrane and receptor organization pathways (Fig. 4D, Supplementary Fig. 3D).

MET and the RTK EGFR share much of the same recycling machinery (56,58), and EGFR is also frequently amplified in MPNST (59). To determine if p53 loss also enhanced EGFR activation, we treated the NF1-MET and NF1-MET;sgP53 cells with EGF in a time course of 5 minutes to 1 day. Strikingly, EGF induced much stronger EGFR phosphorylation in the NF1-MET;sgP53 cells than the parental cell line (Supplementary Fig. 4), similar to the increased MET activation upon HGF treatment. This corresponded with increased and prolonged activation of pERK, pAKT, and pS6. These results suggest an important role for p53 in regulating both MET and EGFR signaling in MPNSTs and may partially explain resistance to EGFR inhibition in MPNSTs (60).

To further examine the effect of p53 expression on MET signaling, we used RPPA to evaluate the response of 98 protein phosphosites to short and extended HGF stimulation. After 2 hours of HGF treatment, numerous phosphosites were significantly upregulated in both the NF1-MET and NF1-P53 cell lines, yet the number of upregulated sites as well as the magnitude of change was higher in the p53-deficient cells (Fig. 4E-F, Table S1). Additionally, phosphorylation of several proteins, including STAT3, JAK2, and B-RAF, were significantly decreased in the NF1-MET cell line after just 2 hours HGF treatment, (Fig. 4E, Table S1) confirming that rapid negative feedback signaling in response to MET activation is p53-dependent. Remarkably, after 48 hours of HGF treatment, only 3/98 phospho-sites were significantly changed in the NF1-MET cells compared to 27/98 sites in the NF1-P53 cells (Fig. 4E-F, Table S1). Notably, STAT3 Y705 phosphorylation, which is sustained by perinuclear MET (61), was among the most significantly increased phosphosites in the p53-deficient cells after 48 hours of HGF exposure (Fig. 4F, Table S1). Collectively, these data suggest that p53 loss moderates MET-addiction by modulating the location, timing, and magnitude of MET effector signaling.

### p53 loss drives mTOR Dependency in MPNSTs

AKT activation was consistently elevated and sustained in p53-deficient MPNST cell lines and is targetable therapeutically, either directly or via its downstream effectors. To determine the scope of AKT/mTOR pathway activation *in vivo*, we assessed the phosphorylation status of mTORC1 and mTORC2 pathway effectors in MPNST tumorgrafts by RPPA. Globally, phosphorylation was significantly increased in NF1-P53 tumors compared to NF1-MET tumors (p-value = 0.0034) (Fig. 5A), with increased activation of 7/12 phosphosites in NF1-P53 tumors (Fig. 5B). AKT and S6RP phosphorylation were significantly increased in the NF1-P53 tumors (Fig. 5B), suggesting increased dependency on the AKT/mTOR pathway. Regardless of p53 status, treatment with the AKT inhibitor, afuresertib, had no effect on the MPNST cell lines, either as a single agent or in combination with trametinib (Supplementary Fig. 5A-B). However, treatment with the mTOR inhibitor, everolimus, significantly decreased viability in the p53-deficient cell lines compared to the p53-intact cells (Fig. 5C). This enhanced inhibition of the p53-deficient cells was observed even though everolimus strongly inhibited downstream pS6RP regardless of p53 status (Fig. 5D). Further, combination therapy of mTOR (everolimus) and MEK (trametinib) inhibition reversed clonal selection for p53 loss (Fig. 5E-F).

**Figure 5.**
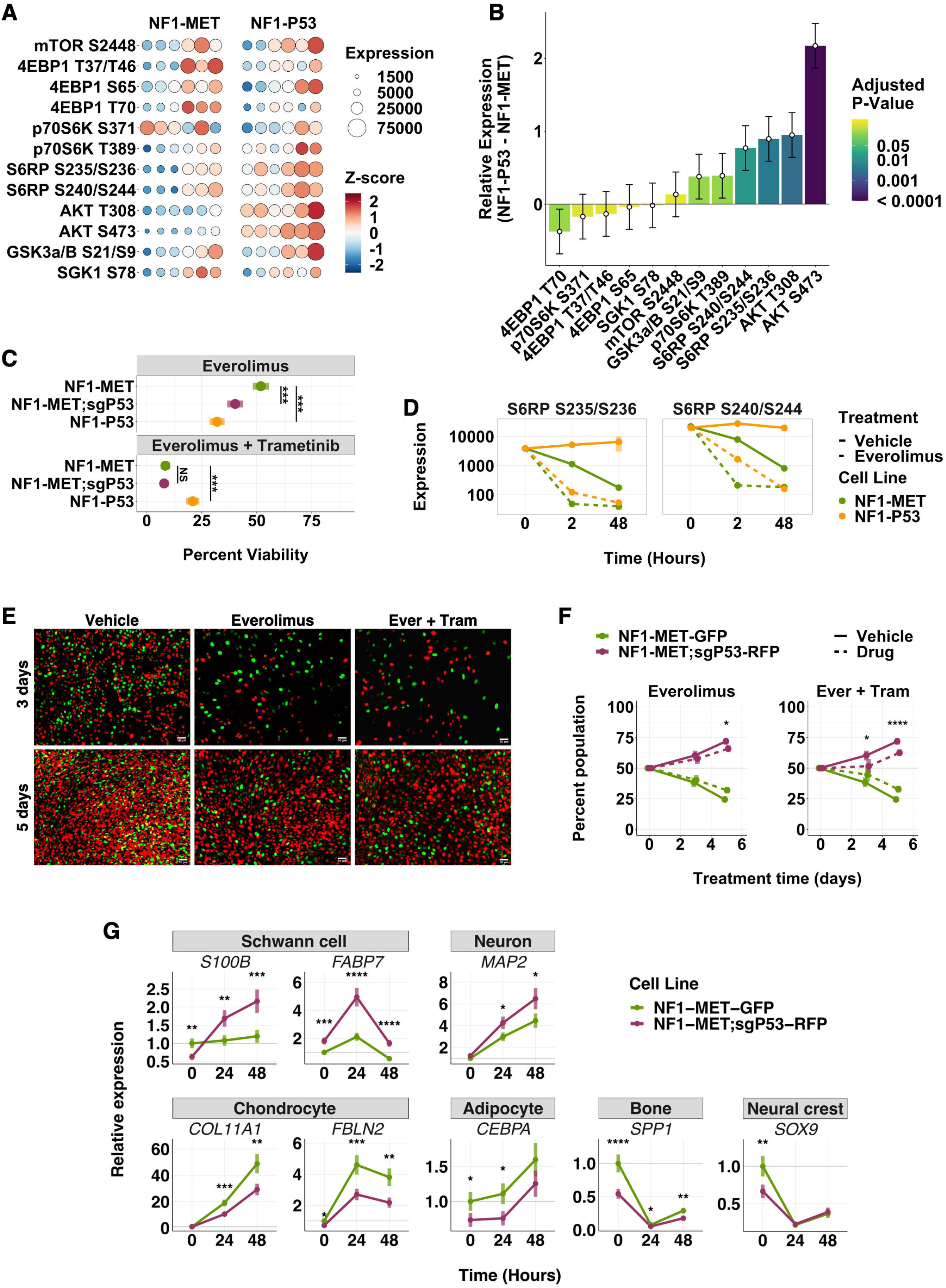
p53 deficiency induces mTOR dependency. (A) mTORC1/mTORC2 phospho protein expression (size) and z - score (color) of NF1-MET and NF1-P53 tumors (6 animals/group). (B) Contrast estimates+/- SE of mTORC1/mTORC2 phospho-protein expression in NF1-P53 tumors compared to NF1-MET tumors. Color indicates P-value. (C) Percent viability of NF1-MET, NF1-MET;sgP53, and NF1-P53 cells after 72 hours of everolimus (20 nM) or combination (everolimus 20 nM, trametinib 40 nM) treatment. (D) Phospho-S6RP expression measured over time by RPPA after vehicle (DMSO) or everolimus (100 nM) treatment. Images (E) and flow cytometry analysis (F) of GFP labeled NF1-MET and RFP labeled NF1-MET;sgP53 cells after 3 and 5 days of treatment with vehicle (DMSO), everolimus (20 nM), or combination everolimus (20 nM) and trametinib (40 nM). (G) Expression (relative to the housekeeping gene PPIA) of cell fate markers by qRT-PCR upon treatment with combination everolimus (20 nM) and trametinib (40 nM). See also Supplemental Figure 6E. * p < 0.05, ** p < 0.01, *** p < 0.001, **** p < 0.0001

As combination mTOR and MEK inhibition was so effective in inhibiting MPNST cell growth, we next asked whether treatment reversed the nuclear localization of MET leading to global downregulation of MAPK and AKT/mTOR signaling. Unexpectedly, combination everolimus and trametinib actually induced ligand-independent MET localization specifically in the NF1-MET;sgP53 cells compared to the parental line (Supplementary Fig. 6A). Moreover, treatment with everolimus or the dual PI3K/mTOR inhibitor BEZ235 induced stronger compensatory ERK and AKT activation in NF1-P53 cells, consistent with increased MET activation (Supplementary Fig. 6B). These data reinforce the role of p53 in regulating MET localization and effector signaling in response to diverse stimuli.

We next tested whether the excessive AKT activation induced by everolimus resulted in p53-independent oncogene induced apoptosis or senescence in p53 null MPNST cells (62,63). Combination everolimus and trametinib treatment did induce apoptosis in the NF1-MET;sgP53 cells compared to the parental cell line, however, the difference was minor (Supplementary Fig. 6C). We also did not observe senescence based on the senescence marker beta-galactosidase after 7 days of drug treatment (Supplementary Fig. 5D).

We evaluated whether differentiation of MPNST cells was impacted by treatment by combined everolimus and trametinib treatment. We observed that mTOR and MEK inhibition in NF1-MET;sgP53 cells significantly increased expression of the Schwann cell markers *S100B* and *FABP7*, yet this increased Schwann cell differentiation was not observed in NF1-MET cells (Fig. 5G). We also examined differentiation marker expression of neuron, chondrocyte, adipocyte, bone, kidney, endothelial, and muscle cells, as well as the multipotency marker *SOX9*, which is expressed in neural crest cells. Combined MEK and mTOR inhibition resulted in increase of the neuronal marker *MAP2* in NF1-MET;sgP53 cells, while chondrocyte and adipocyte differentiation was strongly induced in NF1-MET cells (Fig. 5G). Interestingly, both kidney and muscle markers were strongly induced by treatment in both NF1-MET and NF1-MET;sgP53 line (Supplementary Fig. 6E). Consistent with the induction of multiple differentiation pathways upon mTOR and MEK inhibition, the multipotency marker *SOX9* was decreased in both cell lines (Fig. 5G). These results indicate that lineage differentiation is altered by mTOR and MEK inhibition, with p53-deficient MPNST cells most sensitive to shifts back towards differentiated Schwann cell states.

## Discussion

Resistance to both chemotherapy and targeted kinase inhibition in NF1-related MPNSTs is a daunting clinical challenge. Although MPNSTs harbor complex genomic alterations, copy number gains of RTKs, such as MET, PDGFRα, and EGFR are commonly detected (33,64). Moreover, autocrine MET-HGF signaling has been demonstrated to promote acquired resistance to MEK inhibition (64). Previously, we observed that p53 status impacted the therapeutic response to combined MET and MEK inhibition in *MET*-amplified MPNST tumorgrafts. Even though it is known that p53 is an independent predictor of poor survival and poor response to neoadjuvant chemotherapy in MPNSTs (29,32), how p53 function influences MPNST therapeutic response is not fully understood. We examined whether p53 has a ‘non-canonical’ function and discovered a novel role for p53 in modulating kinome adaptations to targeted therapy. A comprehensive transcriptional and phosphoproteomic analysis revealed multiple mechanisms of resistance, including deregulation of MET stability, localization, and effector activation.

Here we used isogenic models of p53-deficient, *MET*-amplified MPNST cells to identify the distinct adaptive kinome responses to targeted therapies. p53-deficient MPNSTs exhibited increased baseline MET activation suggesting that p53 loss disinhibits amplified MET signaling, an effect that is exaggerated in response to HGF. To determine if p53 is a key driver of clonal selection and drug resistance in MPNSTs, we performed clonal competition assays and discovered that p53-deficient cells had a significant growth advantage in response to MET or MEK inhibition. RNA-seq analysis revealed that p53 regulates expression of several genes involved in MET localization and receptor turnover, resulting in altered signaling kinetics and effector activation. p53 loss resulted in decreased *Cblb* expression, an E3 ubiquitin ligase that targets both MET (54,55) and EGFR (58) for lysosomal degradation. p53 loss also induced *Prkce* expression, which is required for nuclear MET translocation (57), and increased activation of AKT, which is required for nuclear EGFR translocation (65). Collectively, these data suggest a model in which p53 loss causes increased MET localization to the nucleus. In breast and ovarian cancers nuclear MET results in altered effector activation leading to increased calcium signaling (66), PARP activation (53), and YAP dependent transcriptional activation (52) to promote drug resistance and invasion/metastasis. As the majority of MPNSTs demonstrate loss of function of p53, our findings may partially explain the unexpected failure of the EGFR inhibitor erlotinib in a previous clinical trial in MPNST (60). Overall, these data indicate that p53 status should be evaluated as a predictive biomarker of response to RTK inhibition in future MPNST clinical trials.

Even though p53-deficiency promoted resistance to MET and MEK inhibition, we found that p53 loss actually increased sensitivity to mTOR inhibition. Previously, *TP53* loss or mutation has been reported to either decrease (67,68) or increase (62,69,70) sensitivity to mTOR inhibition, depending on the context. Similar to our findings, human rhabdomyosarcoma cells lacking functional p53 are sensitive to rapamycin, while overexpression of p53 induces rapamycin resistance (62). In these studies mechanistically, in the absence of p53, rapamycin induced sustained activation of the ASK1/JNK signaling cascade resulting in persistent c-Jun hyperphosphorylation and subsequent p53-independent apoptosis (63). Restoration of p53 expression resulted in rapamycin resistance by inducing cell cycle arrest instead of JNK/c-Jun-mediated apoptosis (62,63); however, we found no evidence of substantial apoptosis in the context of mTOR inhibition in MPNST. Rather, mTOR and MEK inhibition induced Schwann cell and neuronal differentiation exclusively in p53 null cells in association with increased treatment sensitivity. Normal Schwann cells are capable of undergoing dedifferentiation followed by redifferentiation in cases of peripheral nerve injury to support nerve repair (71). However, these reprogramming pathways are altered in disease states, including NF1, resulting in induction of bone, cartilage, muscle, and adipocyte differentiation pathways (72,73). Induction of differentiation using epigenetic therapies has been proposed as a treatment for sarcomas (74), however our data indicate that kinase inhibitors may also be used to induce differentiation and cell cycle arrest in MPNSTs. Additionally, we show p53 loss plays a paradoxical role in differentiation therapy response, by causing resistance to MET inhibition and increased sensitivity to MEK/mTOR inhibition.

## Conclusion

Our data reveals profound kinome signaling plasticity in MPNST cells and a complex interplay between clonal subpopulations that is influenced by p53. Apart from the well-known tumor suppressor role of p53, we demonstrate that p53-deficiency promotes acquired resistance to targeted kinase inhibition by modulating kinome signaling and MET localization. Moreover, p53-deficiency enhances lineage plasticity which also contributes to kinase inhibitor resistance. Understanding how p53 and other commonly altered genes modulate treatment response is critical for the advancement of precision medicine approaches for MPNST patients.

## Acknowledgements

The authors thank the VARI Vivarium and Transgenics Core and the VARI Optical Imaging Core (confocal microscopy facilities), as well as Zach Madaj for advice on statistical analysis, and Drs Alex Zhong and Payton Stevens for their advice on lentiviral production.

This study was funded in part by grants from the Neurofibromatosis Research Program of the Department of Defense (W81XWH-21-1-0224 and W81XWH-19-1-0537), the Children’s Tumor Foundation Young Investigator Award (CTF-2018-01-009), the Children’s Tumor Foundation (CTF-2015-05-006 and CTF-2019-05-004), NF Michigan, and the Van Andel Institute.

