## Supplemental Figures for "p53 modulates kinase inhibitor resistance and lineage plasticity in NF1-related MPNSTs"

### Slide 1
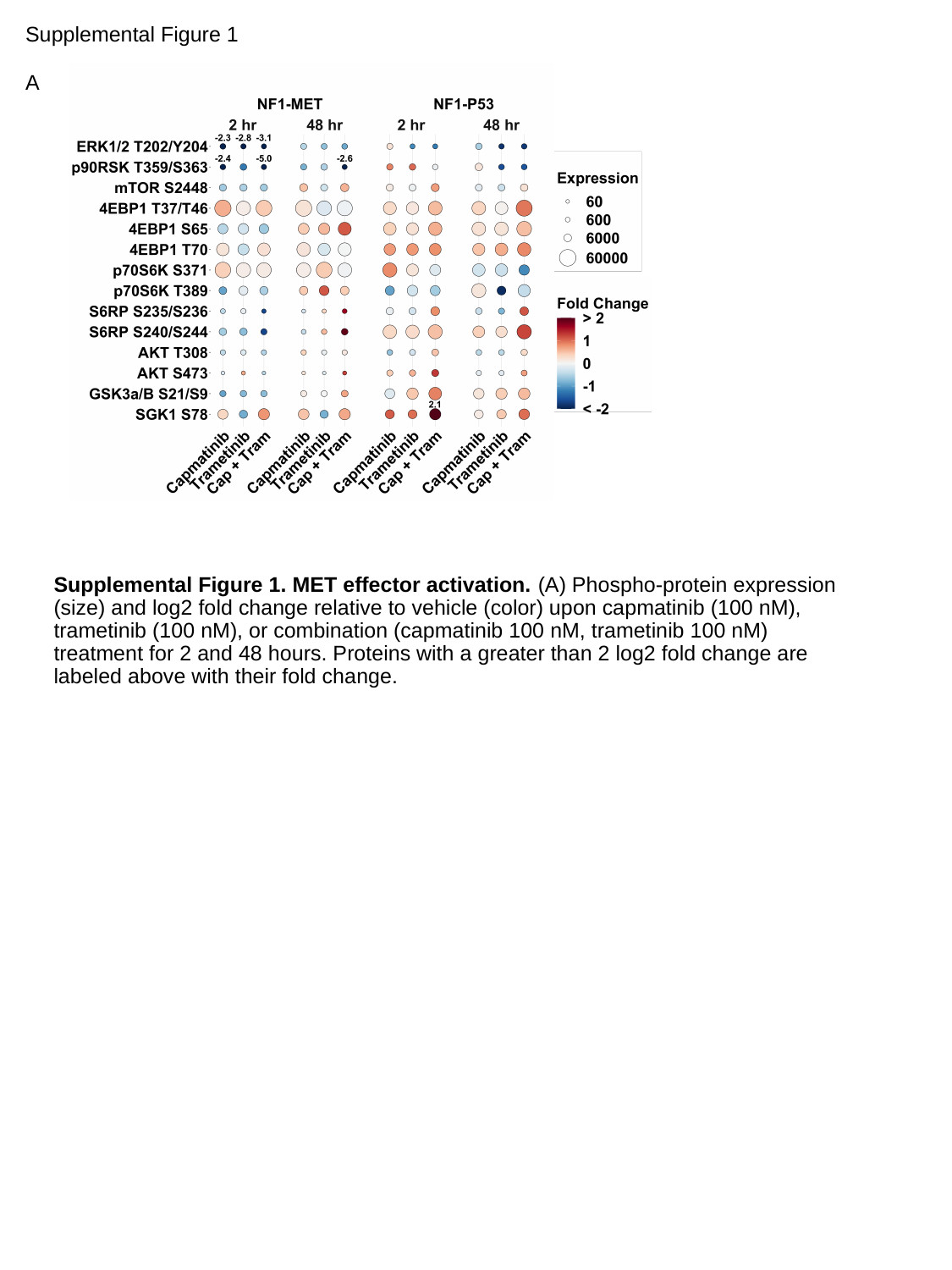

Supplemental Figure 1
A
Supplemental Figure 1. MET effector activation. (A) Phospho-protein expression (size) and log2 fold change relative to vehicle (color) upon capmatinib (100 nM), trametinib (100 nM), or combination (capmatinib 100 nM, trametinib 100 nM) treatment for 2 and 48 hours. Proteins with a greater than 2 log2 fold change are labeled above with their fold change.

### Slide 2
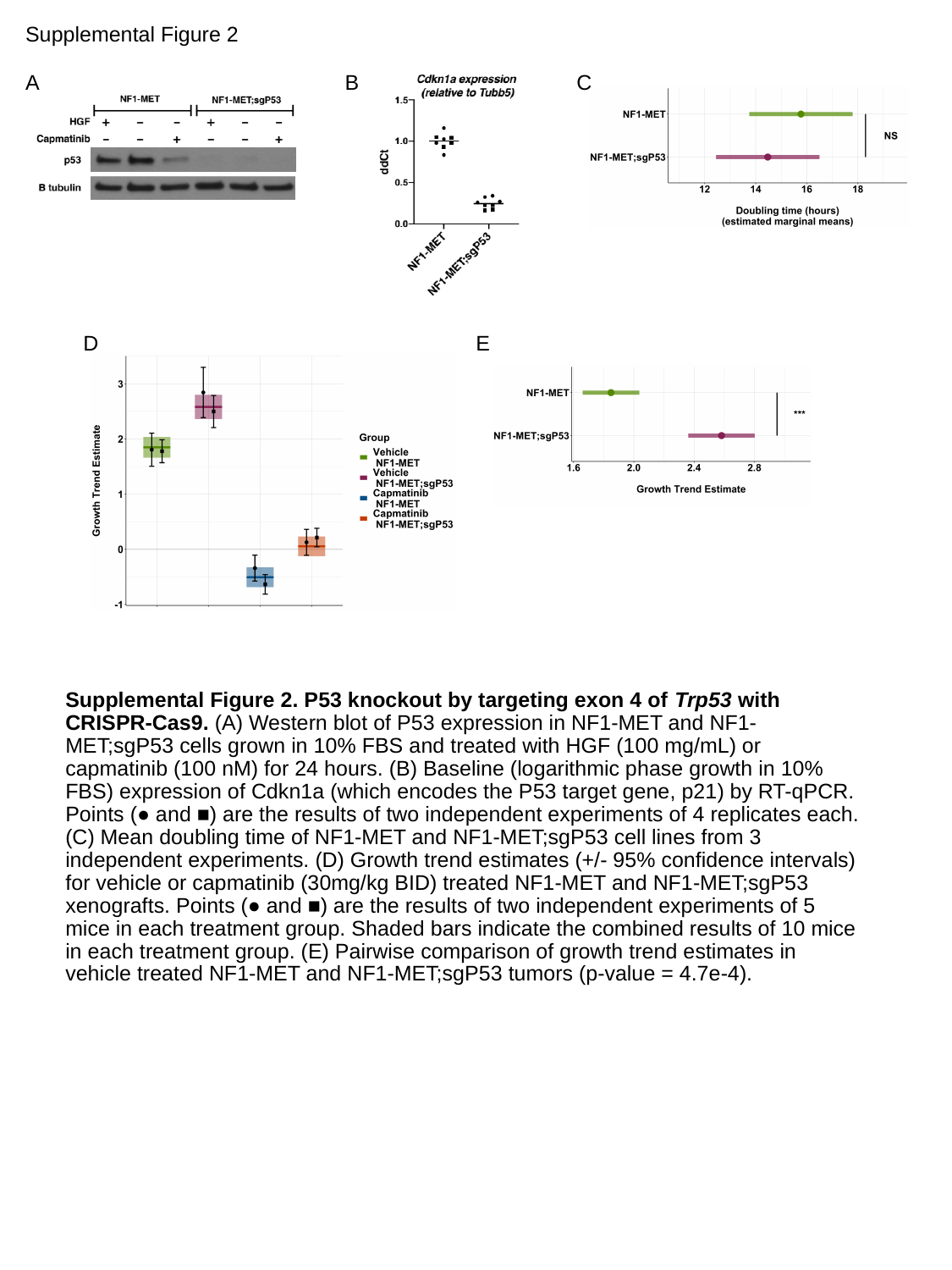

Supplemental Figure 2
C
A
B
E
D
Supplemental Figure 2. P53 knockout by targeting exon 4 of Trp53 with CRISPR-Cas9. (A) Western blot of P53 expression in NF1-MET and NF1-MET;sgP53 cells grown in 10% FBS and treated with HGF (100 mg/mL) or capmatinib (100 nM) for 24 hours. (B) Baseline (logarithmic phase growth in 10% FBS) expression of Cdkn1a (which encodes the P53 target gene, p21) by RT-qPCR. Points (● and ■) are the results of two independent experiments of 4 replicates each. (C) Mean doubling time of NF1-MET and NF1-MET;sgP53 cell lines from 3 independent experiments. (D) Growth trend estimates (+/- 95% confidence intervals) for vehicle or capmatinib (30mg/kg BID) treated NF1-MET and NF1-MET;sgP53 xenografts. Points (● and ■) are the results of two independent experiments of 5 mice in each treatment group. Shaded bars indicate the combined results of 10 mice in each treatment group. (E) Pairwise comparison of growth trend estimates in vehicle treated NF1-MET and NF1-MET;sgP53 tumors (p-value = 4.7e-4).

### Slide 3
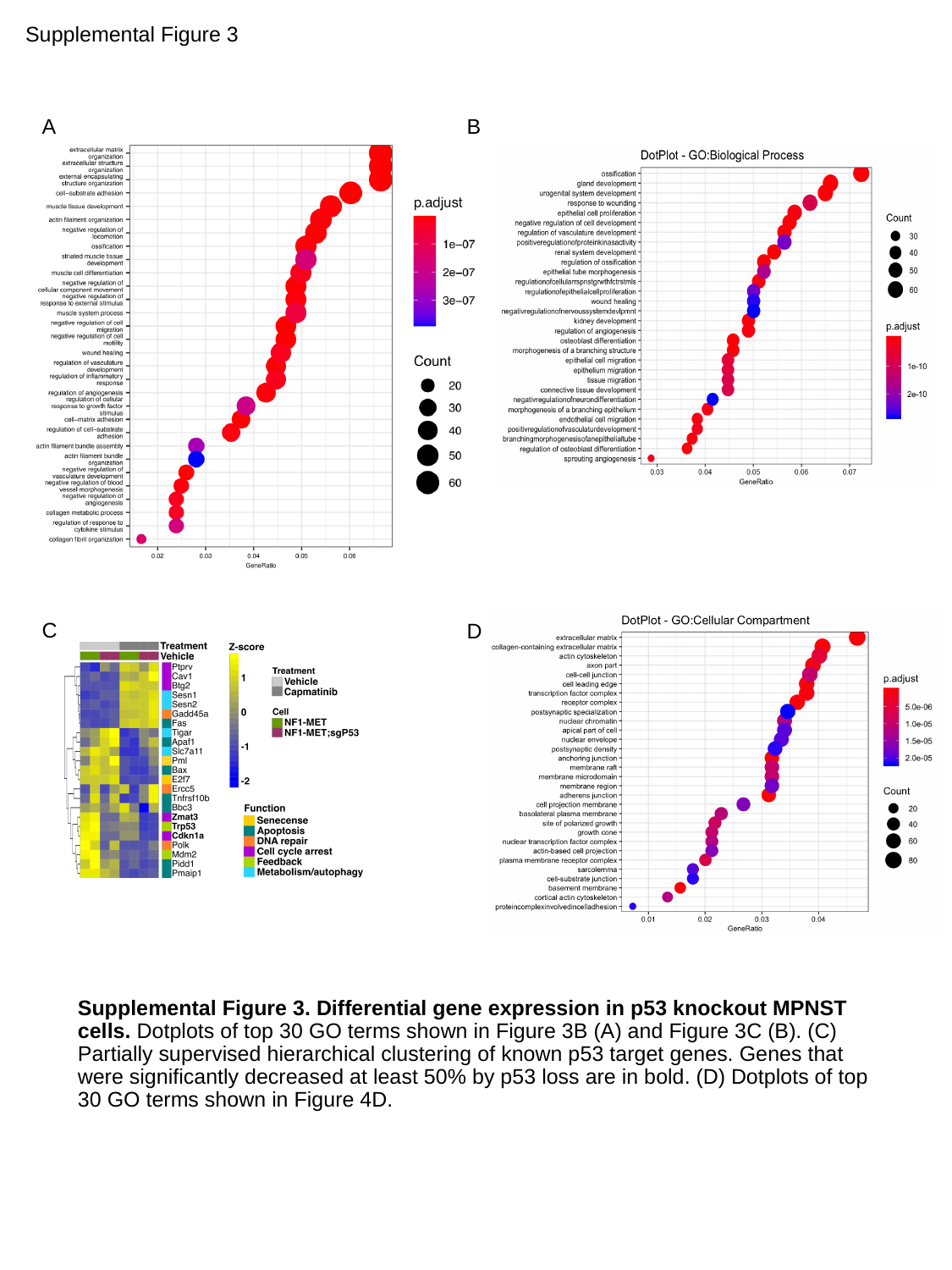

Supplemental Figure 3
A
B
C
D
Supplemental Figure 3. Differential gene expression in p53 knockout MPNST cells. Dotplots of top 30 GO terms shown in Figure 3B (A) and Figure 3C (B). (C) Partially supervised hierarchical clustering of known p53 target genes. Genes that were significantly decreased at least 50% by p53 loss are in bold. (D) Dotplots of top 30 GO terms shown in Figure 4D.

### Slide 4
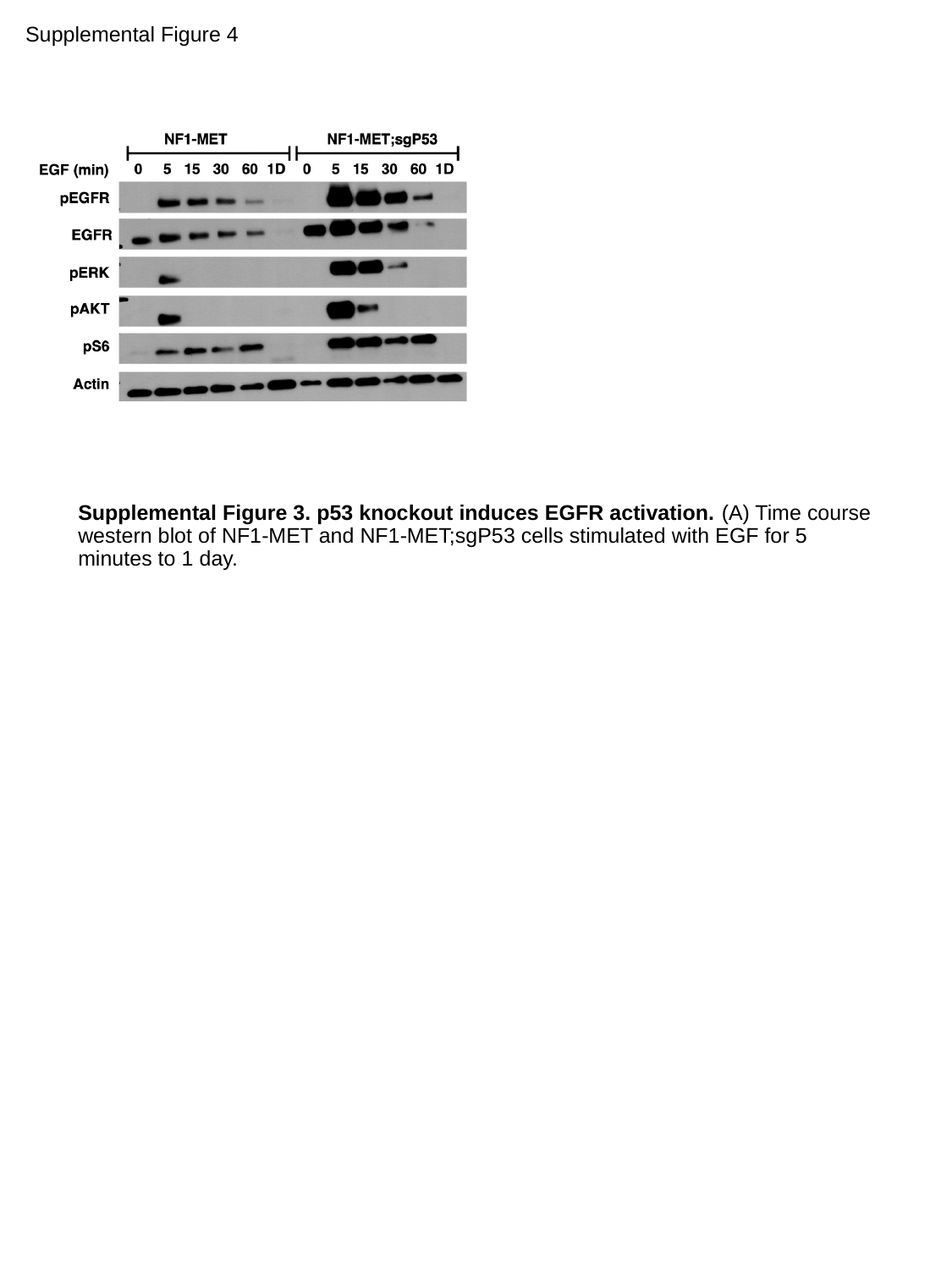

Supplemental Figure 4
Supplemental Figure 3. p53 knockout induces EGFR activation. (A) Time course western blot of NF1-MET and NF1-MET;sgP53 cells stimulated with EGF for 5 minutes to 1 day.

### Slide 5
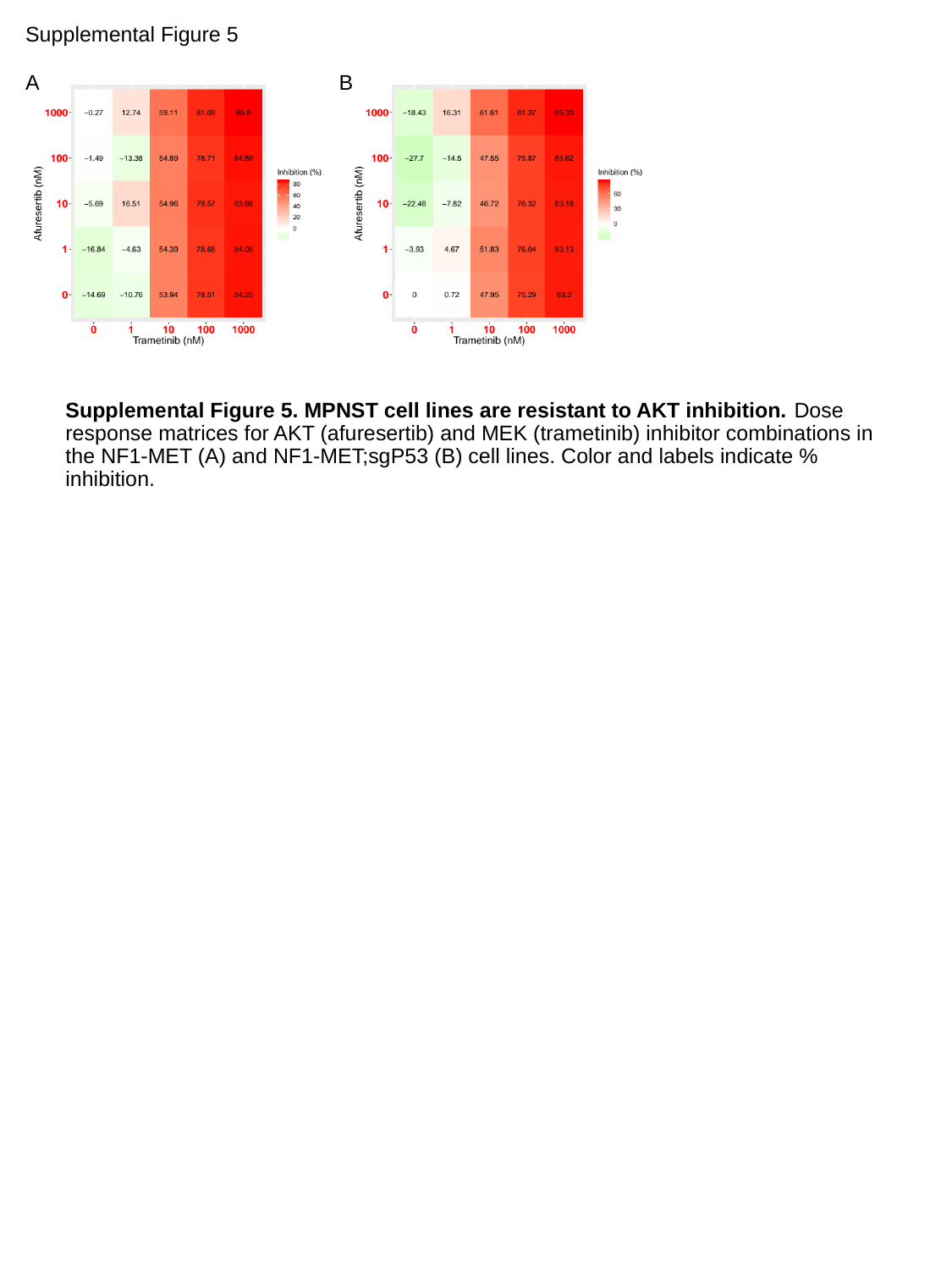

Supplemental Figure 5
A
B
Supplemental Figure 5. MPNST cell lines are resistant to AKT inhibition. Dose response matrices for AKT (afuresertib) and MEK (trametinib) inhibitor combinations in the NF1-MET (A) and NF1-MET;sgP53 (B) cell lines. Color and labels indicate % inhibition.

### Slide 6
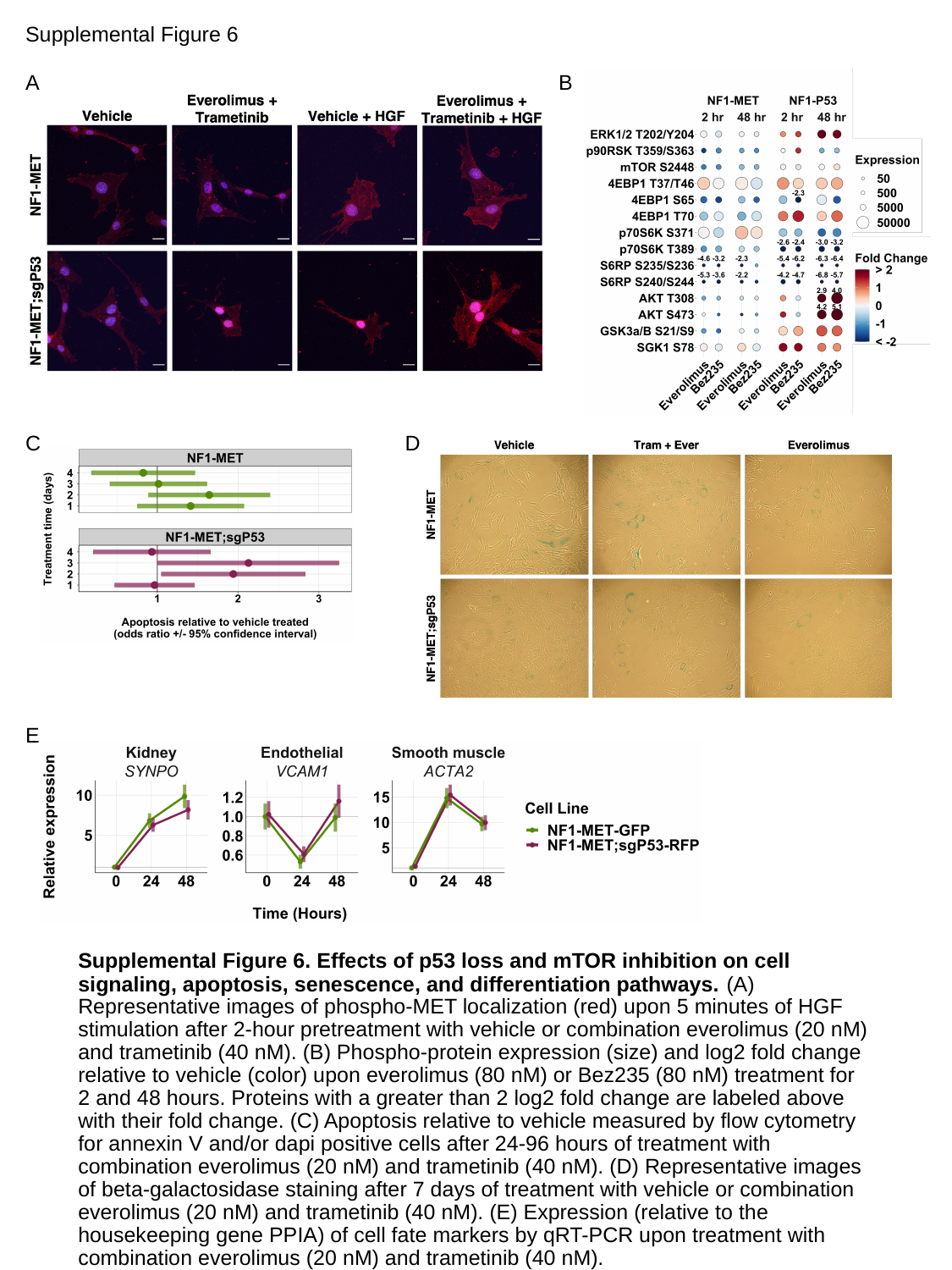

Supplemental Figure 6
B
A
D
C
E
Supplemental Figure 6. Effects of p53 loss and mTOR inhibition on cell signaling, apoptosis, senescence, and differentiation pathways. (A) Representative images of phospho-MET localization (red) upon 5 minutes of HGF stimulation after 2-hour pretreatment with vehicle or combination everolimus (20 nM) and trametinib (40 nM). (B) Phospho-protein expression (size) and log2 fold change relative to vehicle (color) upon everolimus (80 nM) or Bez235 (80 nM) treatment for 2 and 48 hours. Proteins with a greater than 2 log2 fold change are labeled above with their fold change. (C) Apoptosis relative to vehicle measured by flow cytometry for annexin V and/or dapi positive cells after 24-96 hours of treatment with combination everolimus (20 nM) and trametinib (40 nM). (D) Representative images of beta-galactosidase staining after 7 days of treatment with vehicle or combination everolimus (20 nM) and trametinib (40 nM). (E) Expression (relative to the housekeeping gene PPIA) of cell fate markers by qRT-PCR upon treatment with combination everolimus (20 nM) and trametinib (40 nM).

### Slide 7
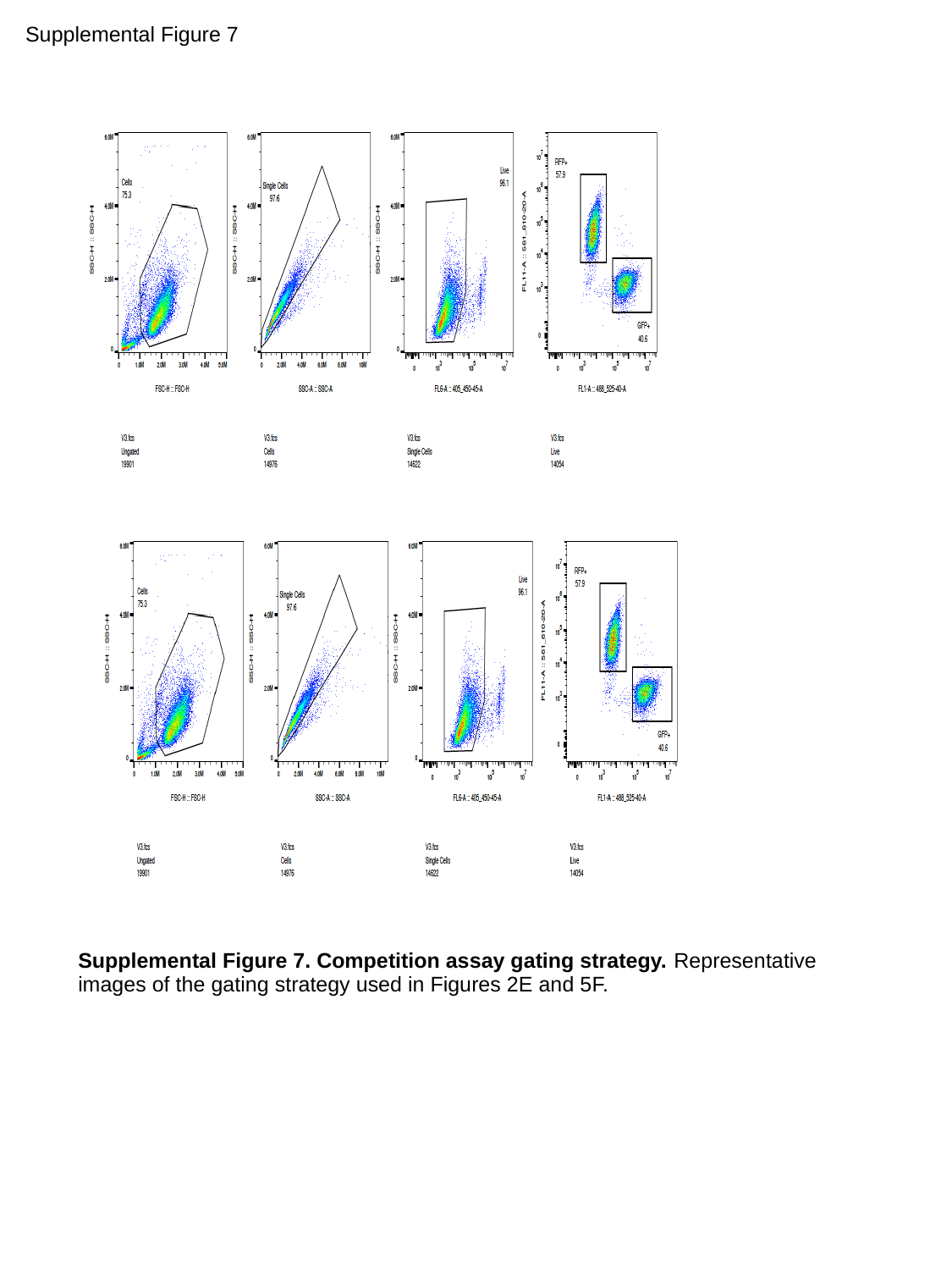

Supplemental Figure 7
Supplemental Figure 7. Competition assay gating strategy. Representative images of the gating strategy used in Figures 2E and 5F.
